# Morphological Attributes of *Glycine max* (Soybean)

**DOI:** 10.1101/2023.03.24.534158

**Authors:** Gazala Bashir

## Abstract

Heavy metal like lead and chromium present in environment in large quantities cause a severe environmental concern. Significant risks to the environment and agriculture are posed by heavy metal contamination of the air, water, and soil as a result of increased urbanisation and industrialization. Heavy metal pollution in soils can harm nearby ecosystems, groundwater, agricultural productivity, and human health because of its persistence and high toxicity. Heavy metals are accumulated in the tissues of plants growing in metal-contaminated soil, which can have a negative impact on the morphology of the plant. The present study was carried out to determine the effect of different concentrations (50ml, 100ml, 150ml and 200ml) of chromium and lead on morphological attributes of three varieties (*JS:335, JS:80-21, JS:75-46*) of soybean(*Glycine max*). The result of the present study showed that the exposure of *Glycine max* to Pb and Cr resulted in a decrease in total length, number of nodes and number of leaves of a plant. The shape of leaves in case of treated plants was also changed from ovoid to heart and oval shapes, at higher concentration of heavy metals. Among the three different varieties (*JS:335, JS:80-21, JS:75-46*) of *Glycine. max*(soybean) *JS:335* showed maximum reduction and the *JS:80-21* showed least reduction in the growth of a plant. Lead treatment proved more toxic than chromium treatment for all the three different varieties (*JS:335, JS:80-21, JS:75-46*) of *Glycine max*.

## 1 Introduction

Heavy metals are elements that are found naturally on Earth. (Vitousek *et al*., 1997). Because they are toxic, persistent, and non-degradable, heavy metals are among the most dangerous contaminants in the aquatic environment. (Proshad *et al*., 2021; Venkateswarlu and Venkatrayulu 2020). Significant hazards to the environment and agriculture are posed by heavy metal contamination of the air, water, and soil as a result of rising urbanisation and industrialisation. Due to its persistence and high toxicity, heavy metal pollution in soils can have detrimental effects on nearby ecosystems, groundwater, agricultural productivity, and human health. This has a significant effect on humanity because agriculture cannot be sustained without healthy soils. (Gunawardana *et al*. 2011). According to numerous studies, the main risk associated with heavy metal contamination of agricultural soils is toxic metal accumulation in crops. Soybean is frequently grown in stressful environments, such as soils with high arsenic concentrations. (Masuda and Goldsmith, 2009; Bundschuh *et al*., 2012;) Soybean (Glycine max) contains protein content of approximately 35–50% of protein depending on its origin. It is a significant source of protein for humans and is also beneficial for animal nutrition. Soybean protein is a good source of many essential amino acids. In addition to these, soybean is a good source of inorganic substances, fatty acids (soybean oil), phytoestrogens, and carbohydrate conjugates. A significant amount of unsaturated fatty acids, B vitamins, and minerals like nitrogen, potassium, magnesium, iron, calcium, and phosphorus enhance their nutritional value. (Rucinski, 2018). Vegetable soybean is regarded as a functional food crop because it is abundant in phytochemicals that are good for humans. (Beckmann *et al*., 1997). The growth of soybean plants exposed to arsenic has shown severe damage: crop yield and root and shoot biomass are both significantly reduced. (Talano *et al*., 2013.heavy metals includes lead and chromium are potentially toxic and are the main sources of environmental pollution. One of the most prevalent and widely dispersed toxic elements in soil is lead (Pb). Plant morphology, growth, and photosynthetic processes are negatively impacted by it. *Spartiana alterniflora* and *Pinus helipensis* seed germination are known to be inhibited by lead. Morzck *et al*., 1982). Pb reduces the dry mass of roots and shoots, germination index, percentage of germination, and root/shoot length. (Amoroso et al 2001). Some plant metabolic processes, including the synthesis of nitrogenous compounds, the metabolism of carbohydrates, and the absorption of water, are inhibited by lead. (Sharma, Dubey, 2005, Hamid *et al*. 2010). In addition to altering soil microorganisms and their activities, which worsened soil fertility, pb contamination in soils also had a direct impact on changes in physiological indices, which further decreased yield. (Majer *et al*., 2002). There are many different ways that Cr toxicity in plants can be seen, from decreased yield to effects on leaf and root growth to inhibition of enzymatic activities and mutagenesis. Chromium adverse effects on plant height and shoot growth (Rout *et al*., 1997).The present study was undertaken to investigate the changes in morphological parameters of soybean under lead and chromium stress.

## 2. Materials and methods

### Plant materials and experimental design

Certified seeds of three varieties of soybean (*Glycine max) (JS:335, JS:80-21, JS:75-46*) were procured from Vindhya herbal testing laboratory and nursery Bhopal, M.P India. Seeds were surface sterilized with 0.1% HgCl_2_ (mercuric chloride) for two minutes. The seeds were washed with distilled water twice and air dried. Soil was divided in 27 parts of 4kg each. 3 parts each were treated with four different concentrations i.e. 50ml, 100ml, 150ml and 200ml of two heavy metals Lead (II) nitrate and Potassium dichromate. Seeds were sown at 2-3cm dept in pot and watering was done once in two days till transfer of plantlets for experiments All experimental pots were placed outdoor for growing after reaching the maturity stage the plants were removed and were used for morphological analysis. Total length, number of nodes, number of leaves and shape of leaves of plants was observed as per study of Kumar *et al*., 2015.

## 3. Results

### 3.1 Morphological studies

### 3.2 Comparative effect of heavy metals on morphology of different varieties of soybean plant

#### 3.2.1 Effect of heavy metal on total length of plant

The effect of different concentrations of heavy metals (lead and chromium) on the total length of treated plants in comparison to control were evaluated after harvesting. The maximum reduction in total length of plant was observed at 200ml treatment in JS:335 followed by JS:75-46 and JS:80-21 under lead stress. Similarly in chromium treatment maximum reduction in total length of treated plant were observed at 200ml treatment in JS:335 followed by JS:75-46, minimum decrease at 200ml was observed in JS:80-21. as shown in figure 1

**Figure 1.**
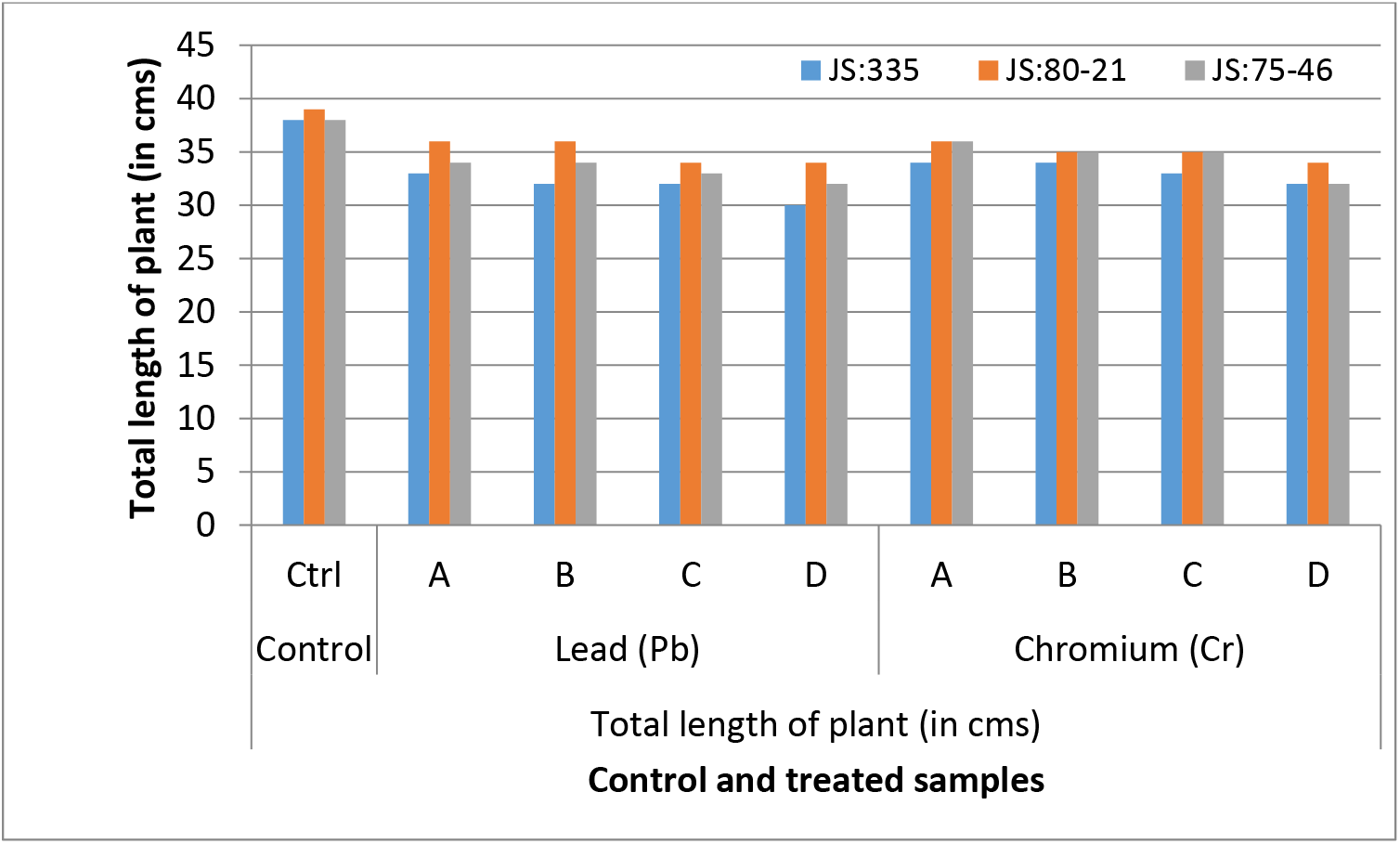
Comparative effect of different concentrations of Lead and Chromium on total length of three varieties (JS: 335, JS: 80-21, JS: 75-46) of *Glycine max* in comparison to control.

#### 3.2.2 Total length of plants (in cms)

#### 3.2.3 Effect of heavy metal on number of nodes of plant

The effect of different concentrations of heavy metals(lead and chromium) on the number of nodes of treated plants in comparison to control were evaluated after harvesting. The maximum reduction in number of nodes of treated plant was observed at 200ml treatment in JS:335 followed by JS:75-46, minimum reduction in JS:80-21 under lead stress. Similarly in chromium treatment maxium reduction in number of nodes of treated plant were observed at 200ml treatment in JS:335 followed by JS:75-46, minium decrease at 200ml was shown in JS:80-21 as shown in figure 2

**Figure 2.**
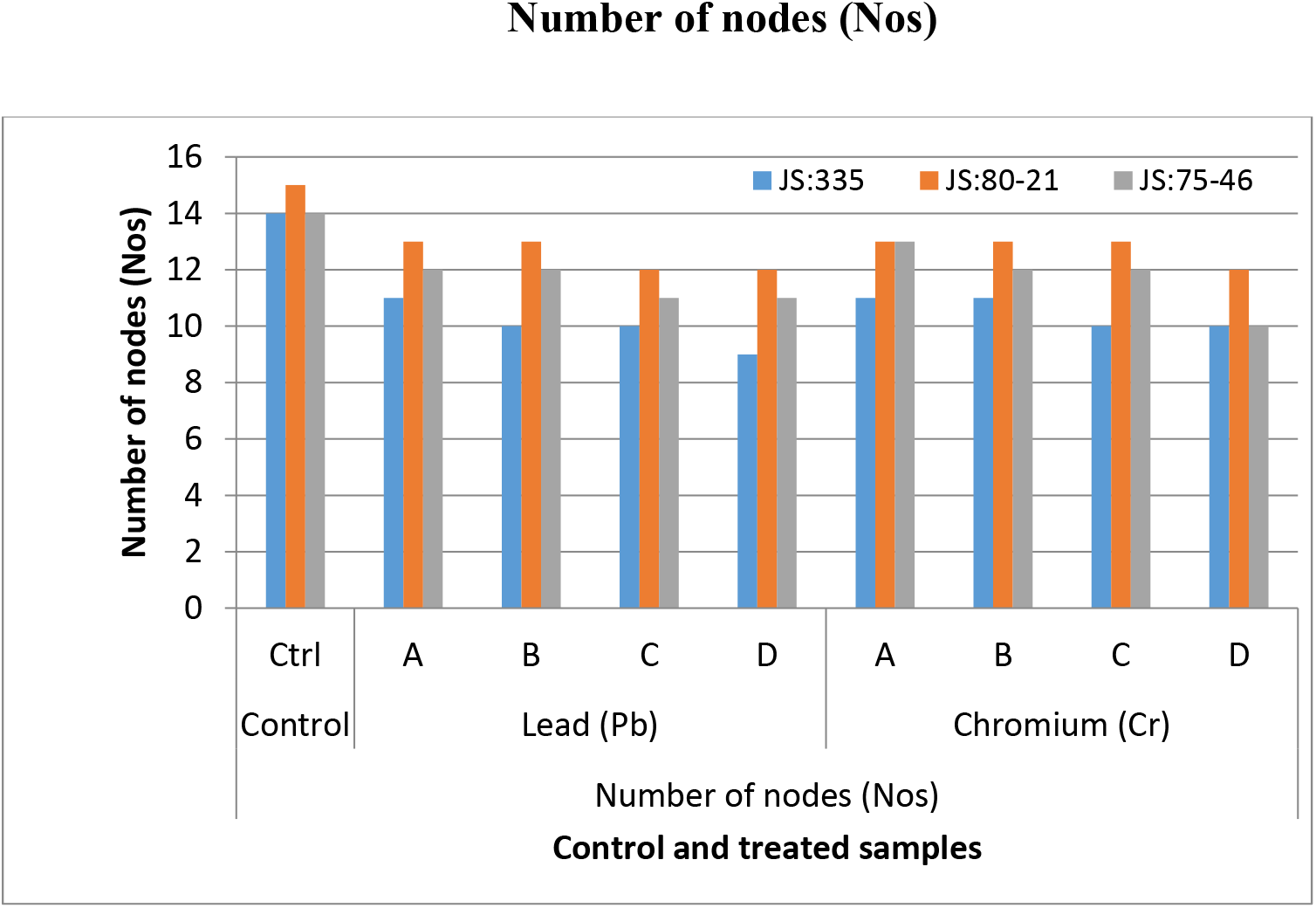
Comparative effect of different concentrations of Lead and Chromium on number of nodes of three varieties (JS: 335, JS: 80-21, JS: 75-46) of *Glycine max* in comparison to control.

#### 3.2.4 Effect of heavy metal on number of leaves of plant

The effect of different concentrations of heavy metals (lead and chromium) on the number of leaves of treated plants in comparison to control were evaluated after harvesting. The maximum reduction in number of leaves of treated plant was observed at 200ml treatment in JS:335 followed by JS:75-46, minimum reduction in JS:80-21 under lead stress. Similarly in chromium treatment maximum reduction in number of leaves of treated plant were observed at 200ml treatment in JS:335 followed by JS:75-46, minimum decrease at 200ml was shown in JS:80-21 as shown in figure 3

**Figure 3.**
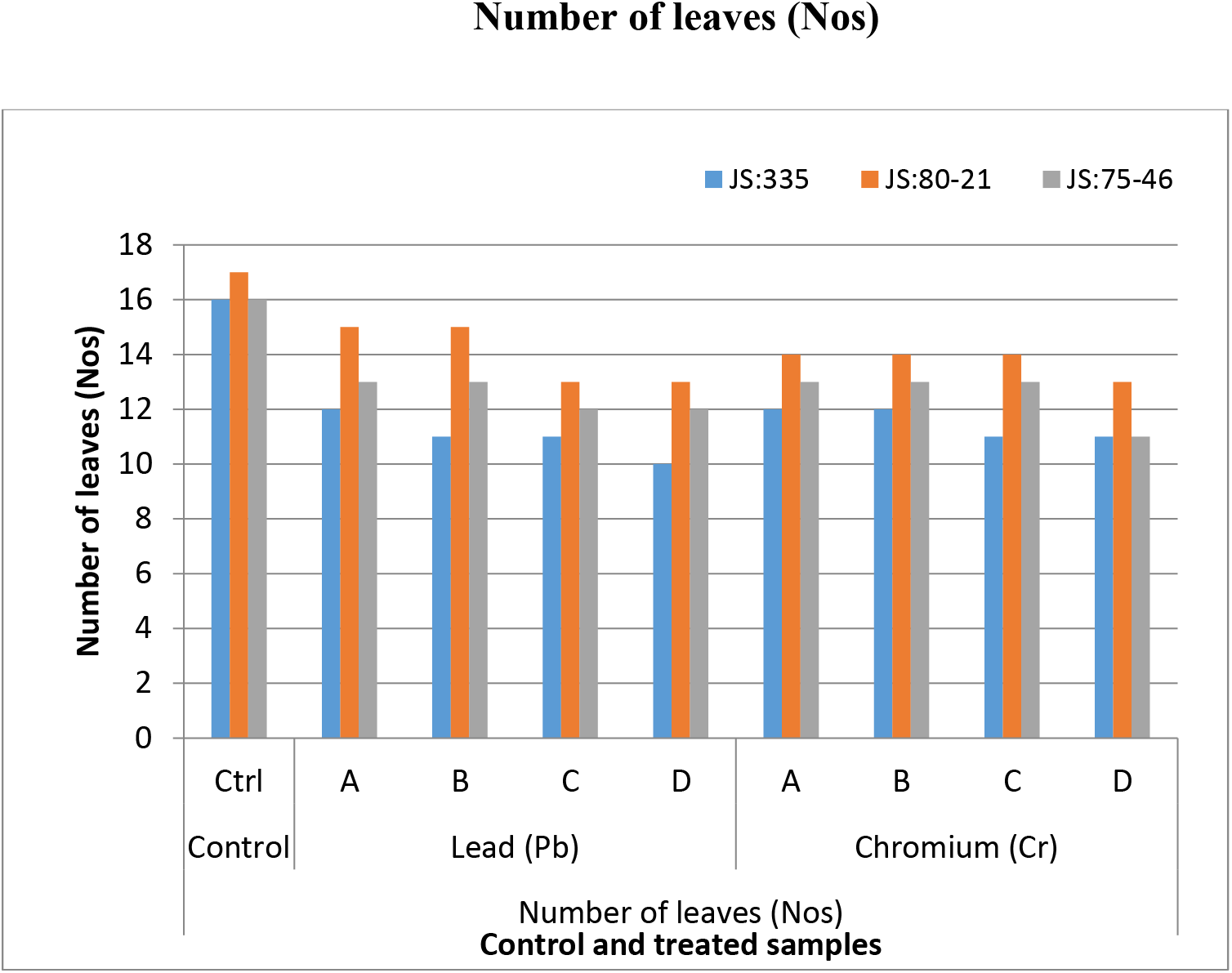
Comparative effect of different concentrations of Lead and Chromium on number of leaves of three varieties (JS: 335, JS: 80-21, JS: 75-46) of *Glycine max* in comparison to control.

#### 3.2.5 Effect of heavy metal on shape leaves

The effect of different concentrations of heavy metals (lead and chromium) on the shape of leaves of treated plants in comparison to control were evaluated after harvesting. Pb and Cr treated plants showed heart and oval shaped leaves over the ovoid shaped leaves in untreated plants. as shown in table 2

**Table 1.**
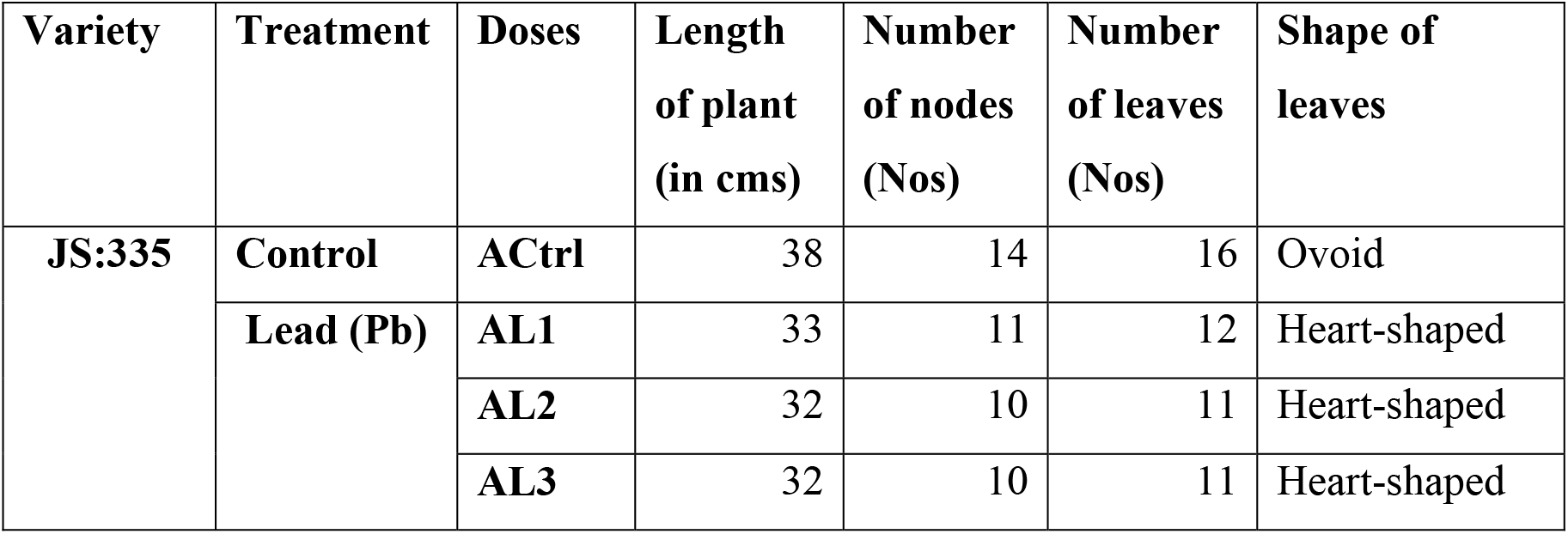
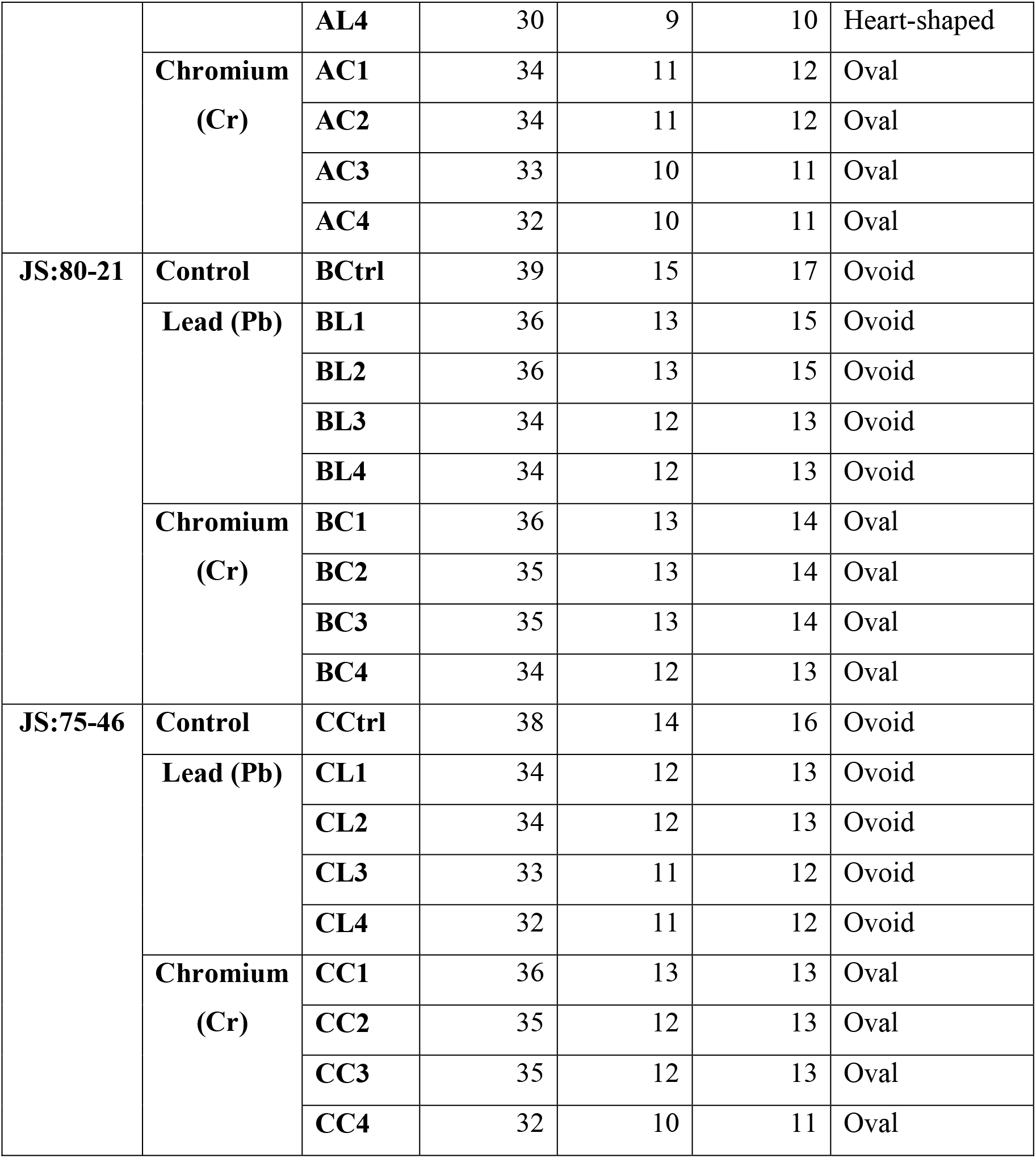
The effect of different concentrations of Heavy metals (Lead and Chromium) on.

**Table 2.** Shape of leaves.

| Variety | Shape of leaves |  |  |  |  |  |  |  |  |
| --- | --- | --- | --- | --- | --- | --- | --- | --- | --- |
|  | Control | Lead (Pb) |  |  |  | Chromium (Cr) |  |  |  |
|  | Ctrl | A | B | C | D | A | B | C | D |
| JS:335 | Ovoid | Heart-shaped | Heart-shaped | Heart-shaped | Heart-shaped | Oval | Oval | Oval | Oval |
| JS:80-21 | Ovoid | Ovoid | Ovoid | Ovoid | Ovoid | Oval | Oval | Oval | Oval |
| JS:75-46 | Ovoid | Ovoid | Ovoid | Ovoid | Ovoid | Oval | Oval | Oval | Oval |

## 4. Discussion

In the present study morphology attributes of the three varieties (*JS:335, JS:80-21, JS:75-46*) of *Glycine max* (soybean) treated with four different concentrations (50ml, 100ml, 150ml and 200ml) of lead (Pb) and chromium (Cr). After the heavy metal treatment morphological characteristics in terms of total length, number of nodes, number of leaves and shape of leaves were observed as per study of Kumar *et al*., 2015.

The data incorporated in Table 1 displays that the results obtained from the present study showed that after Lead (Pb) and chromium (Cr) treatments (50, 100 and 150 and 200 ml), *Glycine max* plant exhibits the reduction in root length, number of nodes, number of leaves at all concentration over control. Also shape of leaves was also deformed in the treated plants. Lead (Pb) proved more toxic than chromium. At Pb treatment, maximum decrease of 21.05 % in total length of plant was observed at 200ml treatment in *JS:335*, followed by 15.8% at 200ml in *JS:75-46* and 12.8 % at 200 ml treatment of heavy metal in *JS:80-21*. The obtained data of number of nodes showed that maximum reduction of 35.71 % was observed at 200ml doses in *JS:335* followed by 21.0% in *JS:75-46* and minimum decrease of 20.0% at 200ml was shown in *JS:80-21*. For number of leaves more reduction was found at 200ml of Pb (37.5%) by *JS:335* and minimum at 200ml (23.52%) by *JS:80-21*.

Similarly in case of Cr treatment maximum decrease of 21.05 % in total length of treated plants was observed at 200ml treatment in *JS:335* followed by 15.7% in JS:75-46, and minimum decrease of 12.8% at 200ml was shown in *JS:80-21*. The obtained data of number of nodes showed that more reduction was found at 200ml (28.57) in *JS:335* and *JS:75-46* and minimum decrease at 200ml (20.0%) in *JS:80-21* plants. The maximum decrease in the number of leaves was also found at 200ml (31.25) in *JS:335* and *JS:75-46* and minimum (23.52%) at 200ml in *JS:80-21*treated plants. Besides the reduction in the total length, no. of nodes and leaves in Pb and Cr treated plants, the shape of leaves also changed. Pb and Cr treated plants showed heart and oval shaped leaves over the ovoid shaped leaves in untreated plants. The investigation revealed significant reduction in total length for G. max plants in Pb treatment than Cr treated plants. A concentration dependent decrease in total length over control was observed in all the three varieties (*JS:335, JS:80-21, JS:75-46*) of *Glycine max*. The results obtained from this study indicate that excess of heavy metal causes a variety of toxicity symptoms in plants, such as reduced growth. According to Eun et al., (2000) the reduction in plant growth during stress is due to low water potential, hampered nutrient uptake and secondary stress such as oxidative stress. Contrary to Cr and Pb treatments, Pb showed more impact on theses parameters. It has been well documented that Cr and Pb, both induce reductions in growth parameters such as fresh mass and dry mass, and shoot and root length, photosynthetic pigments and protein (Panda and Khan, 2003; Tripathi et al., 2012a).

## 5. Conclusion

In conclusion, our results indicated that the exposure of *Glycine max* to Pb and Cr resulted in a decrease in total length, number of nodes and number of leaves. The shape of leaves in case of treated plants was also changed from ovoid to heart and oval shapes, at higher concentration of heavy metals. Among the three varieties (*JS:335, JS:80-21, JS:75-46*) of *G. max, JS:335* showed maximum reduction and the *JS:80-21* showed least less reduction in the growth. Lead proved more toxic than chromium for all the three varieties of *Glycine max*. Still some understandings about the molecular mechanism are needed in order to investigate the process of proper toxic signal pathways of heavy metals exhibited by treated plants.

## Parsed Citations

**Beckman, K. B., & Ames, B. N. (1997). Oxidative decay of DNA. Journal of Biological Chemistry, 272(32), 19633-19636.**

Google Scholar: Author Only Title Only Author and Title

**Bundschuh, J., Litter, M. I., Parvez, F., Román-Ross, G., Nicolli, H. B., Jean, J. S. & Toujaguez, R. (2012). One century of arsenic exposure in Latin America: A review of history and occurrence from 14 countries. Science of the Total Environment, 429, 2-35.**

Google Scholar: Author Only Title Only Author and Title

**Gunawardana, B., Singhal, N., & Johnson, A. (2011). Effects of amendments on copper, cadmium, and lead phytoextraction by**

**Lolium perenne from multiple-metal contaminated solution. International Journal of Phytoremediation, 13(3), 215-232.**

Google Scholar: Author Only Title Only Author and Title

**Hamid, N., Bukhari, N., & Jawaid, F. (2010). Physiological responses of Phaseolus vulgaris to different lead concentrations. Pak J Bot, 42(1), 239-246.**

Google Scholar: Author Only Title Only Author and Title

**Kumar, A., Pandey, A., Aochen, C., & Pattanayak, A. (2015). Evaluation of genetic diversity and interrelationships of agro-morphological characters in soybean (Glycine max) genotypes. Proceedings of the National Academy of Sciences, India Section B: Biological Sciences, 85, 397-405.**

Google Scholar: Author Only Title Only Author and Title

**Majer, B. J., Tscherko, D., Paschke, A., Wennrich, R., Kundi, M., Kandeler, E., & Knasmüller, S. (2002). Effects of heavy metal contamination of soils on micronucleus induction in Tradescantia and on microbial enzyme activities: a comparative investigation. Mutation Research/Genetic Toxicology and Environmental Mutagenesis, 515(1-2), 111-124.**

Google Scholar: Author Only Title Only Author and Title

**Masuda, T., & Goldsmith, P. D. (2009). World soybean production: area harvested, yield, and long-term projections. International food and agribusiness management review, 12(1030-2016-82753), 1-20.**

Google Scholar: Author Only Title Only Author and Title

**Mrozek Jr, E., & Funicelli, N. A. (1982). Effect of zinc and lead on germination of Spartina alterniflora Loisel seeds at various salinities. Environmental and Experimental Botany, 22(1), 23-32.**

Google Scholar: Author Only Title Only Author and Title

**P. Ruciński, Annual Oilseeds and Products Report, FAS, Warsaw, Poland, 2018, https://app.fas.usda.gov.**

**Proshad, R., Islam, S., Tusher, T. R., Zhang, D., Khadka, S., Gao, J., & Kundu, S. (2021). Appraisal of heavy metal toxicity in surface water with human health risk by a novel approach: a study on an urban river in vicinity to industrial areas of Bangladesh. Toxin reviews, 40(4), 803-819.**

Google Scholar: Author Only Title Only Author and Title

**Rout, G. R., Samantaray, S., & Das, P. (1997). Differential chromium tolerance among eight mungbean cultivars grown in nutrient culture. Journal of plant Nutrition, 20(4-5), 473-483.**

Google Scholar: Author Only Title Only Author and Title

**Sharma, P., & Dubey, R. S. (2005). Lead toxicity in plants. Brazilian journal of plant physiology, 17, 35-52.**

Google Scholar: Author Only Title Only Author and Title

**Talano, M. A., Cejas, R. B., González, P. S., & Agostini, E. (2013). Arsenic effect on the model crop symbiosis Bradyrhizobium– soybean. Plant physiology and biochemistry, 63, 8-14.**

Google Scholar: Author Only Title Only Author and Title

**Venkateswarlu, V., & Venkatrayulu, C. (2020). Bioaccumulation of heavy metals in edible marine fish from coastal areas of Nellore, Andhra Pradesh, India. GSC biological and pharmaceutical sciences, 10(1), 018-024.**

Google Scholar: Author Only Title Only Author and Title

**Vitousek, P. M., Mooney, H. A., Lubchenco, J., & Melillo, J. M. (1997). Human domination of Earth’s ecosystems. Science, 277(5325), 494-499.**

Google Scholar: Author Only Title Only Author and Title

